# Living with relatives offsets the harm caused by pathogens in natural populations

**DOI:** 10.1101/2021.02.02.429439

**Authors:** Hanna M. Bensch, Emily O’Connor, Charlie K. Cornwallis

**Author notes:** Corresponding author: Charlie Cornwallis, **Email**.

## Abstract

Living with relatives can be highly beneficial, enhancing reproduction and survival. However, high relatedness can increase susceptibility to pathogens, a phenomenon known as the ‘monoculture effect’. Here we examine if the benefits of living with relatives offsets the harm caused by pathogens, and if this depends if species typically live with kin. Using comparative meta-analysis across plants, animals and bacteria (nspecies= 56), we show that high within-group relatedness increases mortality when pathogens are present. Contrastingly, mortality decreased with relatedness when pathogens were rare, particularly in species that live with kin. Variation in pathogen abundances was lower across groups of relatives, but rates of mortality were more unpredictable. The effects of within-group relatedness were only evident when pathogens were manipulated, suggesting that the harm caused by pathogens is masked by the benefits of living with relatives. These results highlight the importance of kin selection for understanding disease spread in natural populations.

## Introduction

High relatedness between individuals can favour the evolution of cooperative interactions that increase reproductive success and survival (W. D. Hamilton, 1964a, 1964b). For example, it has repeatedly been shown that individuals can pass on their genes indirectly by providing vital resources to relatives and assisting them with tasks that are difficult to do alone, such as caring for offspring (Alexander, 1974; Rubenstein and Abbot, 2017; West et al., 2007). However, living with relatives can also increase susceptibility to pathogens that spread more easily among genetically similar individuals with similar immune defences (Anderson et al., 1986; Hamilton, 1987; Liersch and Schmid-Hempel, 1998; Schmid-Hempel, 1998; Sherman et al., 1998). This phenomenon has been called the ‘monoculture effect’ (Elton, 1958) after it was observed that clonal agricultural crops were highly susceptible to disease outbreaks (Garrett and Mundt, 1999; Tooker and Frank, 2012; Wolfe, 1985; Zhu et al., 2000). More recently it has been established that in natural populations higher genetic similarity between individuals can increase rates of parasitism (Ekroth et al., 2019). However, what is unclear is whether this translates into higher rates of mortality, rendering populations vulnerable to extinction, or whether the benefits of living with relatives are large enough to offset the costs of increased disease risk (Hughes et al., 2002).

Previous research into the effects of relatedness on disease spread have been conducted on an expansive range of species including bacteria, plants and animals. These studies have revealed remarkable variation in how changes in relatedness influence parasitism and mortality. For example, in honeybees, *Apis melifera*, high relatedness among individuals increases the risk of disease and colony death (Tarpy et al., 2013), whereas in Pharoah ants, *Monomorium pharaonis*, high relatedness reduces the abundance of pathogens (Schmidt et al., 2011). Such differences between species may in part be due to how data has been collected. In some studies, relatedness and pathogens have been experimentally manipulated, whereas in other studies relatedness and the abundances of pathogens have simply been measured (‘observational studies’). In observational studies, results can be variable and difficult to interpret because the causality underlying relationships is uncertain (Lively et al., 2014). For instance, in some species, pathogens kill groups of relatives resulting in pathogen abundances being higher in groups of unrelated individuals, the opposite of the true effect (Ben-Ami and Heller, 2005; King et al., 2011; Teacher et al., 2009).

Additionally, variation across species may exist because different mechanisms have evolved to control disease spread among individuals (Loehle, 1995; Romano et al., 2020). In species where relatives interact frequently, selection is predicted to favour the evolution of strategies that mitigate the impacts of pathogens (Loehle, 1995; Romano et al., 2020). Task partitioning, social organisations that limits interactions, and other mechanisms of so called ‘social immunity’ can all prevent disease spread in groups of relatives (Camargo et al., 2007; Liu et al., 2019; Waddington and Hughes, 2010). However, whether species that typically live with kin are better able to cope with pathogens when relatives interact than species that live with non-kin is unclear.

Disease spread through populations may not only been influenced by relatedness increasing average susceptibility to pathogens, but also variation in pathogen abundances across groups. When there are resistant and susceptible genotypes to pathogens, groups of relatives are likely to be either all vulnerable or all protected against pathogens, generating high variation across groups (Boomsma and Ratnieks, 1996; van Baalen and Beekman, 2006). Conversely, groups of unrelated individuals will contain a mix of susceptible and resistant genotypes, leading to more predictable pathogen abundances and rates of mortality across groups. This does, however, depend on the diversity of pathogens that species are exposed to and whether genotype by pathogen interactions occur. When many pathogens are present, groups of related individuals are more likely to be susceptible to at least one pathogen, which can reduce variation in pathogen abundances across groups to a similar level to groups of unrelated individuals (Ganz and Ebert, 2010; van Baalen and Beekman, 2006). While both increases and decreases in variation in rates of parasitism and mortality have been found in specific study species (Ganz and Ebert, 2010; Johnson et al., 2006; Seeley and Tarpy, 2007; Thonhauser et al., 2016), whether relatedness generally increases or decreases variation in pathogen abundances and mortality across groups remains to be established.

Here we use phylogenetic meta-analysis to first examine whether the benefits of interacting with relatives counteract the costs of increased susceptibility to pathogens. Second, we test if the ability to detect such effects is dependent upon experimental manipulations or if they are also evident from observational studies. Third, we ask whether species that live with kin have reduced susceptibility to pathogens when in groups of relatives compared to species that typically live with non-kin. Finally, we investigate whether variation in pathogen abundances and rates of mortality across groups increases with relatedness. The influence of relatedness on mortality and pathogen abundances were quantified by extracting effect sizes (Pearson’s correlation coefficients *r*) from 75 published studies across 56 species. Variation in pathogen abundances and rates of mortality were measured using a standardised effect size of variance that accounts for differences in means, the coefficient of variation ratio (CVR), that was possible to estimate for 25 species.

## Results

### Relatedness and susceptibility to pathogens

Across plants, animals and bacteria relatedness within groups had highly variable effects on rates of mortality and the abundance of pathogens (Figure 1. Posterior mode (PM) and credible interval (CI) of Zr = 0.062 (−0.119, 0.26), pMCMC = 0.402. Table S5). For example, in the grey foam-nest tree frog, *Chiromantis xerampelina*, rates of mortality went up by 22% when average group relatedness increased from 0.25 to 0.5 (Phillip G. Byrne and Whiting, 2011a), whereas in the house mouse, *Mus musculus*, relatedness of groups had little effect on mortality (Thonhauser et al., 2016). Such variation was ubiquitous across all taxonomic groups and was largely independent of phylogenetic history (% of variation explained by phylogeny = 8.195 (0.107, 31.591). Figure 1. Table S5). The relationship between relatedness and mortality was, however, highly dependent on the presence of pathogens (Table S2). Consistent with the idea that groups of relatives are more susceptible to pathogens, mortality increased with relatedness when pathogens were present compared to when they were absent (Difference: PM (CI) = −0.286 (−0.439, −0.117), pMCMC = 0.002. Table S6). The effect of relatedness on the abundance of pathogens in groups was weaker (PM (CI) = 0.096 (−0.1, 0.329), pMCMC = 0.306. Table S6), but not significantly different from the effect of relatedness on mortality (PM (CI) = 0.014 (−0.1, 0.194), pMCMC = 0.512. Table S6).

**Figure 1:**
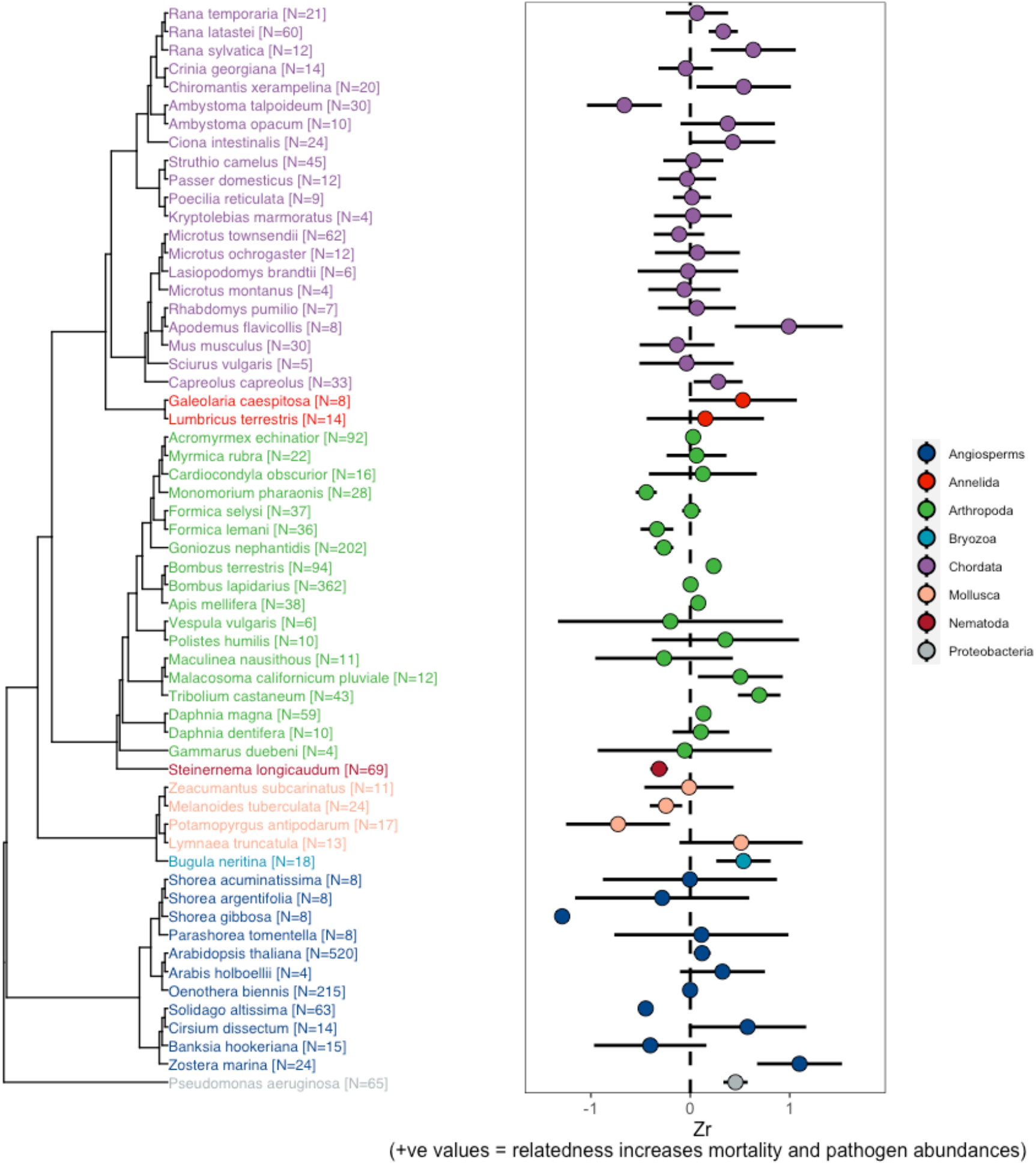
Variation in the effect of relatedness on rates of mortality and pathogen abundance across animals, plants and bacteria. Points represent weighted means for each species and bars are 95% confidence intervals calculated from the sample sizes of the number of groups used studies.

### Experimental studies reveal contrasting effects of relatedness in the presence and absence of pathogens

In support of pathogens having a causal role in increasing mortality in groups of relatives, there were significant differences between studies that experimentally manipulated pathogens compared to observational studies (Table S7). In observational studies, the effect of relatedness on mortality, both in the presence and absence of pathogens, was close to zero (Pathogens present: PM (CI) = 0.059 (−0.095, 0.273), pMCMC = 0.356. Pathogens absent: PM (CI) = 0.1 (−0.114, 0.5), pMCMC = 0.264. Table S7). In contrast, in studies that experimentally manipulated pathogens groups of relatives had higher mortality when pathogens were present relative to when pathogens were absent (Difference: PM (CI) = −0.396 (−0.568, −0.212), pMCMC = 0.001. Table S7). These opposing effects meant that the relationship between relatedness and mortality, both when pathogens were present and when they were absent, did not differ from 0 (Pathogens present: PM (CI) = 0.166 (−0.093, 0.376), pMCMC = 0.148. Pathogens absent: PM (CI) = −0.233 (−0.494, 0.033), pMCMC = 0.108. Figure 2. Table S7). This suggests that the greater susceptibility of groups of relatives to pathogens can be masked by kin selected benefits of living with relatives when pathogens are rare.

**Figure 2:**
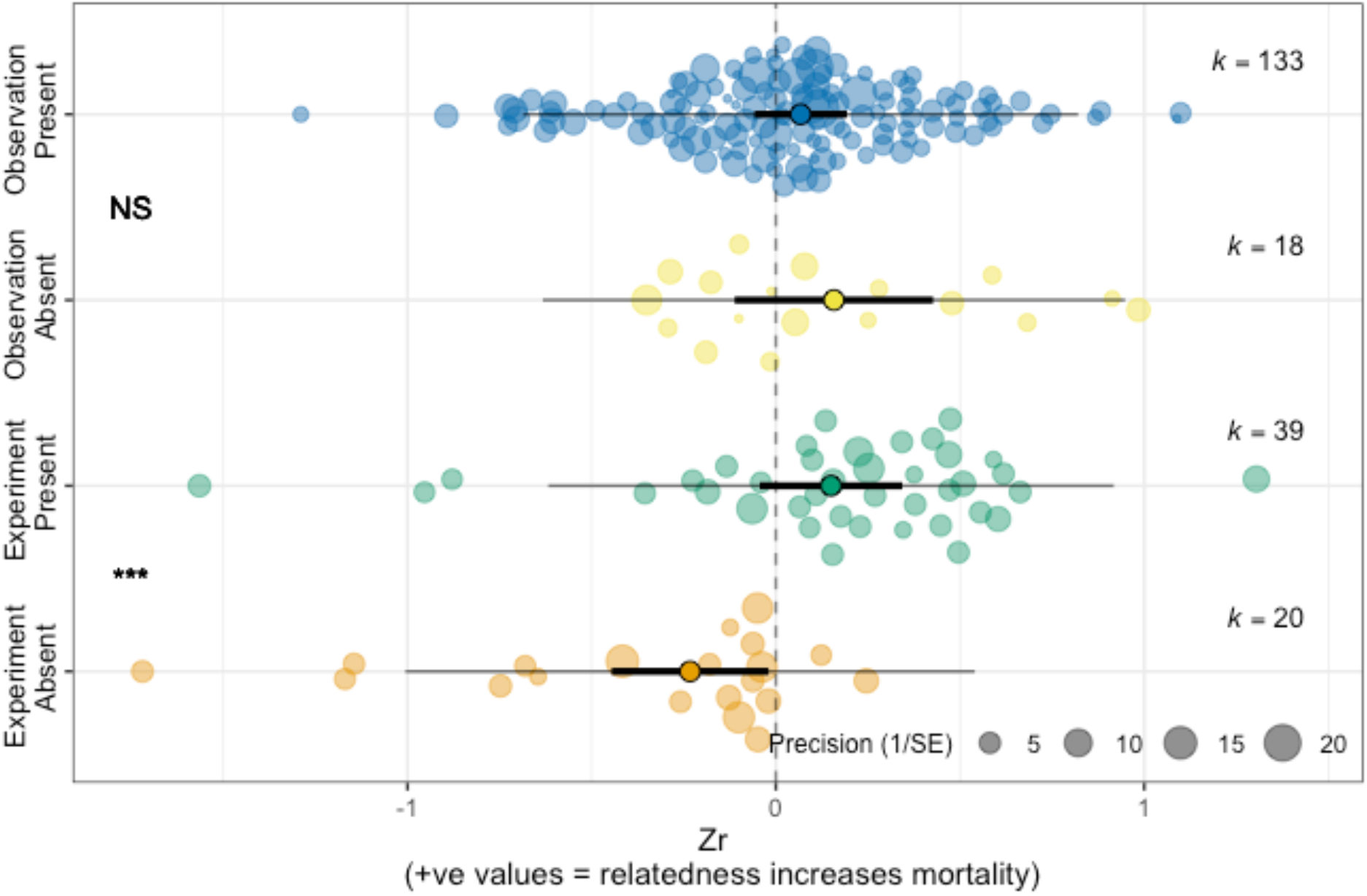
Experimental manipulations are key to detecting the effects of pathogens on groups of relatives. Studies that manipulated pathogen presence demonstrated that groups of relatives had lower mortality when pathogen were absent, but higher rates of mortality when pathogens were present. Points represent means, thick bars are 95% CIs, thin bars are prediction intervals, and k is the number of effect sizes. Each dot is an individual effect size and with size scaled to 1/SE (Orchard plots: Nakagawa et al. (2020)). The statistical difference between comparisons are denoted with symbols (NS = non-significant, * pMCMC <0.05, ** pMCMC<0.01, *** pMCMC<0.001) placed mid-way between comparison groups.

Controlled experiments were less important for detecting the effects of relatedness on mortality. Studies that experimentally manipulated levels of within-group relatedness found that mortality was significantly reduced in highly related groups when pathogen were absent, compared to when then were present (Difference in experimental studies: PM (CI) = −0.191 (−0.418, −0.075), pMCMC = 0.01. Comparable effect sizes were found in observational studies where within-group relatedness was not manipulated (Difference in observational studies: PM (CI) = −0.303 (−0.767, −0.101), pMCMC = 0.022. Table S8).

### Living with kin influences responses to pathogens

Next, we tested whether species that typically live with relatives have evolved mechanisms to limit the negative effects of pathogens when within-group relatedness is high. This was done by comparing the relationship between within-group relatedness and mortality in species that typically associate with relatives (*r* => 0.25 referred to as “kin”) to those that associate with unrelated individuals (*r* <0.25 = referred to as “non-kin”. See methods for details of data used for classifications). When pathogens were present, the effect of relatedness on rates of mortality did not differ between species that live with kin and non-kin (PM (CI) = 0.09 (−0.313, 0.373), pMCMC = 0.79. Table S9). However, when pathogens were absent, higher relatedness reduced mortality in species that live with kin, but increased mortality in species that live with non-kin (PM (CI) = −0.568 (−1.108, 0.024), pMCMC = 0.034. Figure 3. Table S9). For example, in the red flour beetle, *Tribolium castaneum*, and the tube worm, *Galeolaria caespitosa*, that typically interact with non-kin, mortality was two to four times higher when individuals were placed in groups of relatives compared to when individuals were unrelated (Agashe (2009);McLeod and Marshall (2009)). These results show that species that typically associate with non-kin suffer reductions in fitness when placed in groups of relatives, but only in the absence of pathogens. Conversely, species that live with kin had higher fitness in groups of relatives when pathogens were absent, but such benefits dissipated in the presence of pathogens (PM (CI) = −0.333 (−0.534, −0.157), pMCMC = 0.001. Table S9).

**Figure 3:**
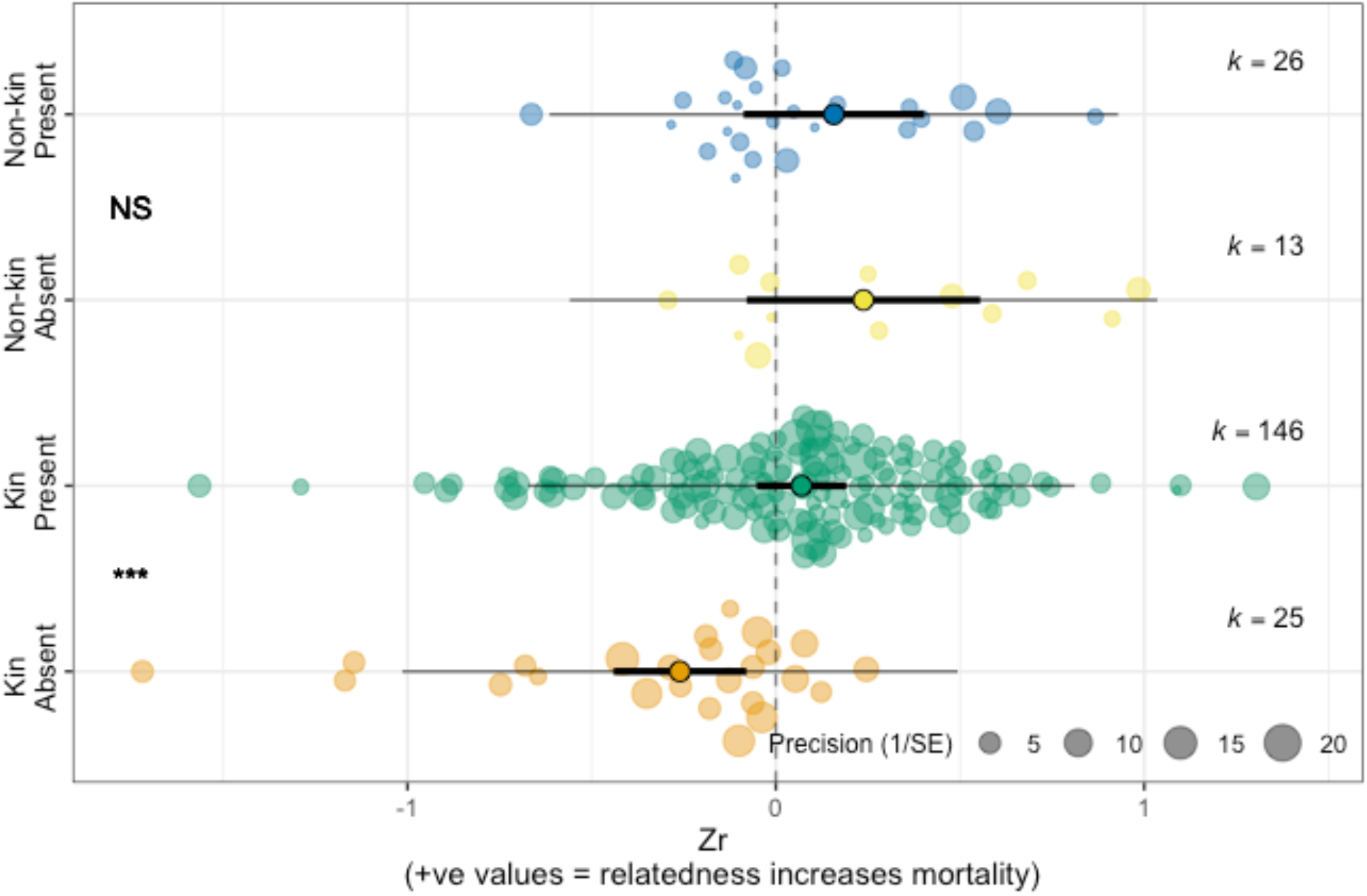
Species that live with kin respond differently to experimental manipulation of pathogen compared to species that live with non-kin. When pathogens were experimentally removed species that live with kin have higher survival, which was reverse when pathogens were present. In contrast, in species that live with non-kin relatedness had no effect when pathogens were absent, but had higher mortality when pathogens were present. Such effects were not evident in observational studies. The components of the orchard plots are the same as in figure 2. The statistical difference between comparisons are denoted with symbols (NS = non-significant, * pMCMC <0.05, ** pMCMC<0.01, *** pMCMC<0.001) placed mid-way between comparison groups.

### Relatedness increases variance in mortality across groups, but decreases variance in pathogens

Variation in rates of mortality and the abundance of pathogens were influenced by relatedness in opposing ways, but this was only evident in experimental studies (Figure 4. Table S10-S13). In observational studies, relatedness within groups had no effect on variance in mortality, either in the presence or absence of pathogens, and did not influence variance in pathogen abundances (mortality pathogens absent: PM (CI) = 0.435 (−0.318, 0.939), pMCMC = 0.322. Mortality pathogens present: PM (CI) = −0.029 (−0.735, 0.648), pMCMC = 0.928. Pathogen abundance: PM (CI) = −0.151 (−0.708, 0.495), pMCMC = 0.832. Figure 4. Table S12). However, in experimental studies higher relatedness was associated with greater variance in mortality in the presence of pathogens (PM (CI) = 0.884 (0.213, 1.413), pMCMC = 0.018. Figure 4. Table S12). The opposite pattern was true for pathogen abundances with there being less variation among groups of relatives compared to group with lower relatedness (PM (CI) = 1.18 (0.557, 1.88), pMCMC = 0.001. Figure 4. Table S12). These results suggest that abundances of pathogens are more consistent across groups of relatives than across groups of unrelated individuals, but whether this translates into mortality is unpredictable.

**Figure 4:**
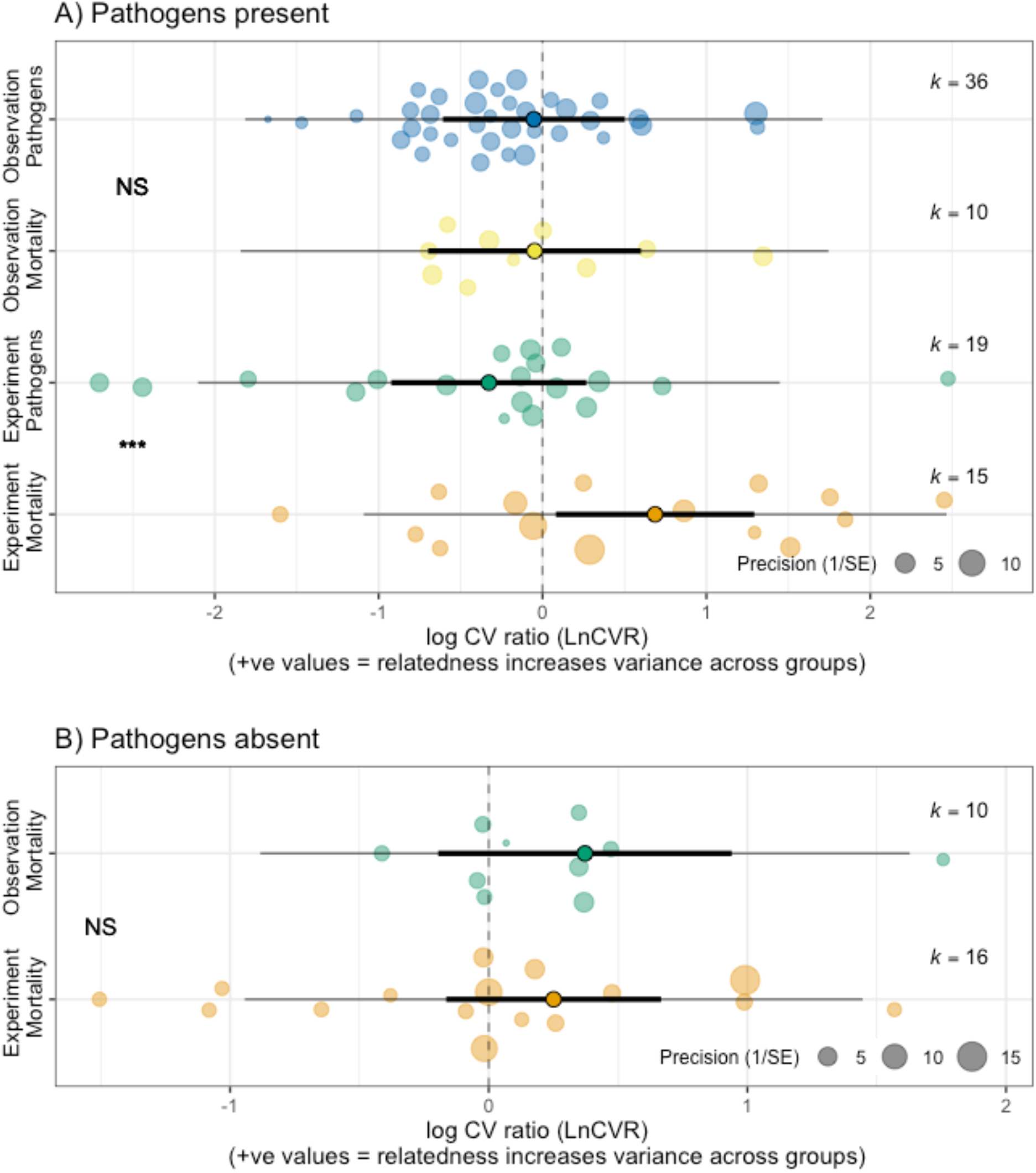
The effect of within-group relatedness on variance in mortality and pathogen abundance. A) In the presence of pathogens, relatedness increased variance in mortality, but decreased variance in pathogen abundance. B) When pathogens were absent, relatedness did not influence variance in mortality. The components of the orchard plots are the same as in figure 2. The statistical difference between comparisons are denoted with symbols (NS = non-significant, * pMCMC <0.05, ** pMCMC<0.01, *** pMCMC<0.001) placed mid-way between comparison groups.

## Discussion

Previous research has shown that high relatedness among individuals can aid the spread of disease and increase mortality (Anderson et al., 1986; Liersch and Schmid-Hempel, 1998; Sherman et al., 1998). Our results support this idea, but show that the detrimental effects of pathogens can be counteracted by the benefits of living with relatives. Such contrasting effects of relatedness mean that experimental manipulations of pathogens are crucial to understanding the impact of diseases on natural populations and accurately quantifying the net benefits of living with relatives. Individuals of species that associate with kin appear to benefit from being in groups of relatives more than species that typically interact with non-kin, but only when pathogens are rare. This suggests that the evolution of social groups containing highly related individuals may be more likely to occur in environments where pathogens are scare. The relatedness structure across groups also appears to influence how variable the effects of pathogens are. While pathogens appear to spread more easily across groups of relatives, rates of mortality are more variable making it difficult to predict the effect of pathogens in more genetically structured populations.

Why have mechanisms not evolved to reduce the susceptibility of groups of relatives to pathogens? The benefits to living with kin should generate selection for increased resistance or tolerance to disease spread among relatives (Loehle, 1995; Romano et al., 2020). However, even in species where individuals obligately live with kin there was little evidence that they were able to reduce the harm caused by pathogens. One explanation is that species that live in highly related groups may increase genetic diversity within groups when occupying areas where pathogens are more prevalent (Schmid-Hempel, 1998; Sherman and Morton, 1988). For example, increases in mating promiscuity under higher disease risk can lower relatedness among offspring recruited to groups (Busch et al., 2004; Singh et al., 2015; Soper et al., 2014). Such responses can reduce disease spread, but also weaken selection for adaptations that limit pathogen spread amongst highly related individuals.

Alternative defence mechanisms against pathogens have also been documented in species that live with kin (Loehle, 1995; Romano et al., 2020). Social distancing through reduced group size, or changing the organisational structure of groups, can reduce pathogen transmission (Liu et al., 2019). For instance, in leaf cutter ants, *Acromyrmex spp*, young workers are less likely to have encountered pathogens and they perform tasks only within the inner colony. As workers age they move to more peripheral tasks where pathogen exposure is higher, such as foraging and tending waste heaps, and at this point they stop entering the inner colony (Camargo et al., 2007; Waddington and Hughes, 2010). The relative costs of decreasing pathogen susceptibility by reducing relatedness versus other mechanisms remains unclear, but may provide insight into why some species are able to live with relatives even when the risks of disease are high.

In the presence of pathogens, high relatedness was associated with higher and more variable rates of mortality. In contrast, relatedness had little effect on variation in pathogen load, and if anything reduced variation across groups. Such differences may arise because pathogen abundances are only weakly related to virulence across pathogens (Leggett et al., 2012). Furthermore, when genotypes are equally susceptible to pathogens, but vary in their ability to clear infections, then within-group relatedness can influence mortality rates without strongly affecting variation in pathogen loads (Best et al., 2008; Howick and Lazzaro, 2014; Koskela et al., 2002).

The effects of relatedness on variation and mean mortality rates were only evident in experiments. There are a number of possible, non-mutually exclusive, explanations for this. First, it is possible that observational studies fail to capture the true effect of pathogens because of sampling biases due to pathogens removing groups of relatives (Ben-Ami and Heller, 2005; King et al., 2011; Teacher et al., 2009). Second, the diversity and abundance of pathogens species face may differ between experimental and observational studies, which are often conducted on wild populations (Table S1). Although experiments often reported that pathogens were manipulated in biologically realistic ways, it is possible that pathogen abundances are generally higher in experiments leading to larger effect sizes. In addition, experiments generally only manipulated single pathogens whereas observational studies on natural populations often involved communities of pathogens. Low pathogen diversity is predicted to increase variation across groups of relatives (Boomsma and Ratnieks, 1996; van Baalen and Beekman, 2006). The lack of an effect of relatedness on variance in mortality in observational studies may therefore be because there was higher pathogen diversity. In our dataset, there was only one experimental study that manipulated multiple pathogens. In *Daphnia magna* it was found that variance in parasitism was higher in groups of relatives (‘clonal’ versus ‘polyclonal’ populations), but this diminished as the number of pathogens increased (Ganz and Ebert, 2010). This suggests that where pathogen diversity is high, groups of relatives become increasingly susceptible to pathogens, reducing variance across groups (Boomsma and Ratnieks, 1996; Parsche and Lattorff, 2018; van Baalen and Beekman, 2006).

The interaction between kin selected benefits and pathogen induced mortality has important implications for the ecological distributions of species. In some lineages, such as birds, cooperative species that live in families have been found to inhabit areas that are hot and dry (Cornwallis et al., 2017; Jetz and Rubenstein, 2011; Lukas and Clutton-Brock, 2017). This has been explained through the benefits of cooperative offspring care being higher where environmental conditions are challenging for independent reproduction (Emlen, 1982). However, it is also possible that the costs imposed by pathogens of living with relatives may be lower in such environments (Campbell-Lendrum et al., 2015). Escaping from pathogens may be an important selective force determining the ecological niches of species with different levels of sociality and relatedness within groups (Altizer et al., 2011). Although the effects of relatedness on pathogen spread have been investigated in a diverse range of species, experimental data on species with different social systems that inhabit different environments is limited. We hope that our results stimulate further research by highlighting the importance of manipulating pathogen presence, abundance and diversity across species with different social systems and ecological niches. This appears crucial to understanding the impact of pathogens on natural populations.

## Materials and Methods

### Literature searches

A systematic literature review was performed to identify studies that have examined the relationship between within-group relatedness and rates of mortality or the abundance of pathogens. One challenge with locating relevant literature was that some studies use the term relatedness while others use the term genetic diversity. Genetic diversity in turn encompassed studies that have examined within-individual genetic diversity (e.g. heterozygosity), as well as genetic diversity within groups. The aims of our study only relate to variation in genetic diversity within groups (relatedness). All studies where estimates of within-group genetic diversity were potentially influenced by within-individual genetic diversity were excluded (see below).

The literature search was performed using the Web of Science (WoS) including articles published up to the 27th July 2020. Searches were restricted to articles in English and the WoS categories were restricted to Beh. Sci, Ecology, Biology, Evol. Ecology, mult. Sci, Genetics h., Biodiv. Cosv., Entomology, Zoology, as a preliminary study (Bensch MSc Thesis) showed these categories to be the ones of interest. Web of science searchers included the following combinations of terms in the topic field: (((“genetic diversity” OR “genetic variability” OR “genetic diversities”) AND parasite*) OR ((“genetic diversity” OR “genetic variability” OR “genetic diversities”) AND disease*) OR ((“genetic diversity” OR “genetic variability” OR “genetic diversities”) AND pathogen*) OR ((“genetic diversity” OR “genetic variability” OR “genetic diversities”) AND survival) OR ((“genetic diversity” OR “genetic variability” OR “genetic diversities”) AND mortality) OR (relatedness AND pathogen*) OR (relatedness AND disease*) OR (relatedness AND parasite*) OR (relatedness AND mortality) OR (relatedness AND survival*) OR “monoculture effect” OR “Monoculture effect”) AND (population* OR group* OR colony). The search yielded a total of 4616 returns, 4615 after removing a duplicate.

To aid finding relevant papers, abstracts were downloaded and imported into R for text-analysis using the quanteda-package (Benoit et al., 2018). The frequency of words in each abstract was calculated and used to create a relevance score according to the number of words with positive and negative interest for this study. The following words had positive associations (listed in order of priority): “genetic”, “diversity”, “diversities”, “variation”, “relatedness”, “related”, “unrelatedness”, “unrelated”, “diverse”, “parasite”, “ectoparasite”, “ectoparasites”, “parasites”, “pathogen”, “pathogenic”, “pathogens”, “disease”, “diseases”, “diseased”, “mortality”, “survival”, “resistance”, “infection”, “infections”, “prevalence”, “tolerance”, “transmission”, “population”, “group”, “colony”, “groups”, “colonies”, “populations”. The following words had negative associations: “human”, “humans”, “hospital”, “cancer”, “hiv”, “patients”. Papers were sorted according to their relevance scores and then manually screened to examine whether they contained data that could be used to calculate an effect size of relatedness and mortality and/or pathogen abundance.

We stopped screening after 2102 papers as number of new papers selected for in-depth screening decreased to less than 1% per 100 references (Figure S1). In addition to WoS searches, reference lists of key-studies and the papers from which we extracted effect sizes were screened for relevant primary literature. PDF files of articles selected based on abstract screening were downloaded for full text, in-depth examination. A preferred Reporting Items for Meta-Analyses diagram (PRISMA Moher et al., 2009) of the literature screening process is shown in figure S2). In total our dataset consisted of 210 effect sizes from 75 studies and 56 species ((Abdi et al., 2020; Agashe, 2009; J. David Aguirre and Marshall, 2012; J. D. Aguirre and Marshall, 2012; Altermatt and Ebert, 2008; Anton et al., 2007; Baer and Schmid-Hempel, 2001; Baer and Schmid-Hempel, 1999; Ben-Ami and Heller, 2005; Bensch and Cornwallis, 2017; Bichet et al., 2015; Byrne and Robert, 2000; Phillip G. Byrne and Whiting, 2011b; Cook-Patton et al., 2017, 2011; Crutsinger, 2006; Gregory M Crutsinger et al., 2008; Gregory M. Crutsinger et al., 2008; Dagan et al., 2017, 2013; de Morais et al., 2020; Desai and Currie, 2015; de Vere et al., 2009; Dobelmann et al., 2017; Ellison et al., 2011; Field et al., 2007; Franklin et al., 2012; Fraser et al., 2010; Gamfeldt and Källström, 2007; Ganz and Ebert, 2010; Gardner et al., 2007; He and Lamont, 2010; Hoggard et al., 2013; Hughes and Stachowicz, 2004; Hughes and Boomsma, 2006, 2004; Johansson et al., 2007; Johnson et al., 2006; Kapranas et al., 2016; Keeney et al., 2009; King et al., 2011; Kotowska et al., 2010; Lambin and Krebs, 1993; Liersch and Schmid-Hempel, 1998; Mattila et al., 2012; McLeod and Marshall, 2009; Mott et al., 2019; Neumann and Moritz, 2000; Page Jr et al., 1995; Parker et al., 2010; Parsche and Lattorff, 2018; Pearman and Garner, 2005; Reber et al., 2008; Robinson et al., 2013; Schmidt et al., 2011; Seeley and Tarpy, 2007; Sera and Gaines, 1994; Shykoff and Schmid-Hempel, 1991; Siemens and Roy, 2005; Solazzo et al., 2014; Strauss et al., 2017; Tarpy, 2003; Tarpy and Seeley, 2006; Tarpy et al., 2013; Teacher et al., 2009; Thonhauser et al., 2016; Trouvé et al., 2003; Ugelvig et al., 2010; van Houte et al., 2016; Vanpé et al., 2009; Walls and Blaustein, 1994; L. A. Wauters et al., 1994; Weyrauch and Grubb, 2006; Winternitz et al., 2014; Woyciechowski and Król, 2001)).

### Overview of study design and inclusion criteria

Studies were included if they presented data on the abundance/presence of pathogens and relatedness for four or more groups. Relatedness was estimated from breeding designs, pedigrees and using molecular markers. A group was defined as three or more individuals as it has been shown to be sufficient for group-level defences (Hughes and Stachowicz, 2004). That said, only three studies used groups with three individuals (4%) with over 93% of studies using groups with five or more individuals. Some studies manipulated levels of relatedness by experimentally creating groups (referred to as ‘experimental’), whereas other studies measured relatedness on already established groups (referred to as ‘observational’). The presence and abundance of pathogens was also experimentally manipulated in some studies (referred to as ‘experimental’) whereas in others pathogens were measured without any manipulations (referred to as ‘observational’).

Studies on plants were included that examined the effect of pathogens and herbivores, as it has previously been argued that herbivory is equivalent to parasitism (see Price 1980 & Siemens and Roy 2005 for discussion of herbivores as pathogens:(Price, 1980; Siemens and Roy, 2005)). One study was included from unpublished data collected by the authors on ostriches, *Struthio camelus* (Tables S19). Studies were excluded if they were on domestic species or where there was the potential for within-individual genetic diversity, including inbreeding, to influence estimates of within-group relatedness. In some studies, inbreeding was not explicit but potentially possible (see SI table S1). We tested the sensitivity of our results to any potential inbreeding effects, by removing these effect sizes and repeating our analyses (see verification analyses Tables S14 & S15). If data of interest were missing in the text or figures, authors were contacted for supplementary data or clarification. If authors did not respond within three months the effect sizes were excluded. If studies provided multiple measures of pathogen load and/or mortality, separate effect sizes were extracted. Where studies presented abundances of pathogens separately as well as total abundance of pathogens the total was used.

### Effect size calculation on the effect of relatedness on rates of mortality and pathogen abundances

The relationship between within-group relatedness and mortality and/or pathogen abundance was analysed by studies either by comparing groups with high and low relatedness (relatedness as a categorical variable), or by analysing variation in average within-group relatedness as a continuous variable. Information from both types of study were used to calculate a standardised effect size of the correlation between within-group relatedness and mortality/pathogen abundance: Pearson’s correlation coefficient, *r*. The statistical tests presented in studies were converted to *r* using the online Meta-analysis calculator (Morris, 2019) and the R package ‘esc’(Lüdecke, 2019). Measures of r were transformed to Zr using ‘escalc’ function in the R package metafor (Viechtbauer, 2010).

In some studies, it was not possible to obtain effect sizes directly from the statistics reported in studies, but *r* could be calculated from data presented in the text and/or figures in two ways. First, in studies where groups with high and low relatedness were compared, means ± SD of mortality or pathogen abundances were used to calculate *r*. Second, in studies where descriptive statistics (e.g. means ± SD) were reported for multiple groups that varied in relatedness, we conducted our own Pearson’s correlations in R (see R script “EffectSizeCalculations” and Table S2 column “Effect size Rscript reference”). In such cases, variation in measures of relatedness, mortality, and pathogen abundances were included by creating distributions from descriptive statistics that were sampled to create 1000 datasets. For each of these 1000 datasets *r* was calculated and an average taken across the 1000 datasets.

### Variance effect size calculations

The effect of relatedness on variance in mortality and pathogen abundances was calculated using the natural logarithm of the ratio between the coefficient of variation from groups with high and low relatedness (LnCVR: (Nakagawa et al., 2015)). LnCVR provides a standardised measure of differences in the variability of two groups accounting for differences in the means between groups. LnCVR was used because estimates of variation increased with the mean (figure S3). LnCVR was calculated from studies that presented means and SDs (converted to SD if studies presented SEs or CIs) across groups when relatedness was low and high. This provides a standardised measure of the effect of relatedness on variability across groups, not within groups (SDs were of the means of groups not individuals).

### Data on study characteristics

For each effect size extracted from each study we collected information on: 1) whether pathogens were present or absent; 2) whether pathogens were experimentally manipulated; 3) whether relatedness was experimentally manipulated; 4) the method used for measuring relatedness (pedigree or molecular markers); and 5) whether pathogen abundance or mortality were measured (where survival estimates were presented the sign of the effect size was reversed). If there was no mention of pathogens in the paper then pathogens were assumed to be present when studies were conducted in the field and absent if conducted in the laboratory.

### Data on species characteristics

For all species in our dataset we searched for whether they typically associate with kin (‘kin’) or not (‘non-kin’) during the life stage that effect sizes were measured. Species were categorised as kin if they lived in groups in which r was estimated to be equivalent to 0.25 or higher and ‘non-kin’ if they live in groups in which relatedness was estimated to be lower than 0.25 (Table S4). Three sources of information were used to estimate levels of relatedness among individuals: 1) estimates of relatedness acquired either directly from molecular genetic analyses or records of groups of individuals with known relatedness; 2) information on the mating system; and 3) typical dispersal patterns as low dispersal from groups increases relatedness. The relevant information was collected using google scholar by including each species name combined with “genetic diversity”, “relatedness” and “group” as search terms to collect measures of within-group relatedness; “mating system” and “paternity” for information on mating system; and “dispersal” and “philopatry” for information on dispersal. The categorization of each species as kin or non-kin along with evidence and the list of literature to support these classifications can be found in Table S4 ((Abdi et al., 2020; J. David Aguirre and Marshall, 2012; J. D. Aguirre and Marshall, 2012; Amiri et al., 2017; Anton et al., 2007; Arnaud and E, 1999; Avise and Tatarenkov, 2015; Barrett et al., 2005; Bee, 2007; Beermann et al., 2015; Ben-Ami and Heller, 2005; Bryja et al., 2008; Byrne and Robert, 2000; Phillip G. Byrne and Whiting, 2011b; Chapuisat et al., 2004; Croshaw et al., 2009; Gregory M Crutsinger et al., 2008; Dagan et al., 2013; Dean et al., 2006; de Morais et al., 2020; de Vere, 2007; de Vere et al., 2009; Dobelmann et al., 2017; Edenbrow and Croft, 2012; Farentinos, 1972; Ficetola et al., 2010; Field et al., 2007; Franklin et al., 2012; Fredensborg et al., 2005; Gamfeldt and Källström, 2007; Gardner et al., 2007; Getz et al., 1993; Goldberg et al., 2013; Goulson et al., 2002; Goymann, 2009; Graham, 1941; Griffin, 2012; Haag et al., 2002; He et al., 2004; Head and Yu, 2004; Heppleston, 1972; Heske and Ostfeld, 1990; Hoffmann et al., 2003; Hoggard et al., 2009; Hughes and Stachowicz, 2004; Johnson, 2007; Johnson et al., 2006; Kapranas et al., 2016; Kawamura et al., 1991; Keeney et al., 2009; Kelly et al., 1999; Keough, 1989; Keough and Chernoff, 1987; Kimwele and Graves, 2003; King et al., 2011; Kozakiewicz et al., 2009; König, 1993; Lambin and Krebs, 1991; Laurila and Seppa, 1998; Lepais et al., 2010; Liker et al., 2009; Liu et al., 2013; Mackiewicz et al., 2006; McLeod and Marshall, 2009; Meling-lópez and Ibarra-Obando, 1999; Myers et al., 2011; Oettler and Schrempf, 2016; Osváth-Ferencz et al., 2017; Pai and Bernasconi, 2007; Pietrzak et al., 2010; Platt et al., 2010; Reusch et al., 1999; Rice et al., 2009; Rock et al., 2007; Russell et al., 2004; Schmid-Hempel and Crozier, 1999; Schmid-Hempel and Schmid-Hempel, 2000; Schmidt et al., 2011, 2016; Schradin et al., 2010; Schrempf et al., 2006; Sepp and Walin, 1996; Seppä et al., 2009; Shapiro and Dewsbury, 1986; Siemens and Roy, 2005; Simeonovska-Nikolova, 2007; Solomon et al., 2004; Stürup et al., 2014; Sutcliffe, 2010; Svane and Havenhand, 1993; Tarpy, 2003; Tatarenkov et al., 2007; Thonhauser et al., 2016; Trouvae et al., 2003; Vanpé et al., 2009; Verrell and Krenz, 1998; Walck et al., 2001; Waldman, 1982; Walls and Blaustein, 1994; Wauters et al., 1990; Wauters and Dhondt, 1992; L. Wauters et al., 1994; Zenner et al., 2014)).

### Data limitations

Our dataset highlighted that there are several key variables where data is limited and where further empirical work would be extremely useful. In particular, information on the following is currently limited: 1) species that typically live with non-kin (*r*: kin=41, non-kin=15. *LnCVR:* kin=18, non-kin=7); 2) studies that quantify the effect of relatedness on rates of mortality in the *absence* of pathogens, particularly under natural conditions. Out of 75 studies, pathogens were excluded in 16 laboratory studies and no studies tried to explicitly exclude pathogens under field conditions. For *LnCVR*, pathogens were only excluded in 7 laboratory studies out of a total of 32 studies; and 3) variation across groups in rates of mortality and pathogen abundance (out of 210 mean effect sizes, variance could only be examined in 106).

### Statistical analysis

#### General techniques

Data were analysed using Bayesian Phylogenetic Mixed Models (BPMM) with Markov chain Monte Carlo (MCMC) estimation and Gaussian error distributions in R package MCMCglmm (Hadfield, 2010). Data points were weighted by the inverse sampling variance associated with each of the effect size using the “mev” term in MCMCglmm.

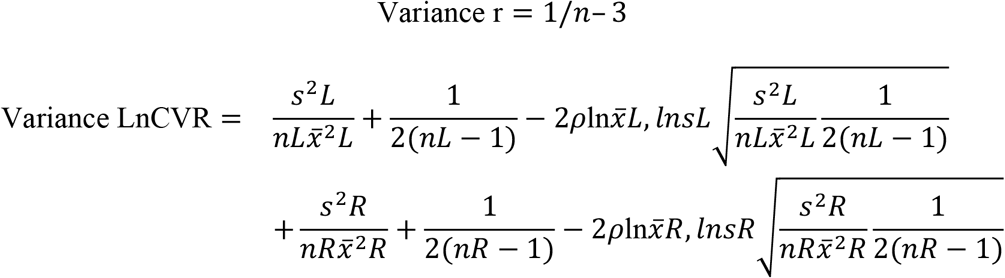

where n corresponds to the number of groups, L and H are groups with low and high relatedness respectively. Unfortunately, the difference in relatedness between low and high relatedness treatments could not be included as a moderator in analyses as exact estimates of relatedness were not always given (e.g. monogamous versus polyandrous breeders) or comparable across studies (e.g. estimates of relatedness from molecular markers do not always equate to relatedness estimates from pedigrees/breeding designs).

The non-independence of data arising from multiple effect sizes per study was modelled by including study as a random effect. In one study (Reber et al., 2008) there were three relatedness treatment groups (low, intermediate and high) allowing effect sizes between low and intermediate, and high and intermediate to be calculated. However, we excluded comparisons with the intermediate treatment to avoid non-independence of effect sizes within studies (Noble et al., 2017). The non-independence of data arising from shared ancestry was modelled by including a phylogenetic variance-covariance matrix of species relationships as a random effect.

The phylogenetic variance-covariance matrix was created from hierarchical taxonomic classifications using the ‘as.phylo’ function in the R package ‘ape’(see Figure 1). We also attempted to create a phylogeny using information from the open tree of life (Rees and Cranston, 2017) using the R package ‘rotl’ (Michonneau et al., 2016). This produced a tree that was extremely similar, but several mollusc species were missing and we therefore used the tree created from taxonomy in all analyses. Branch lengths were estimated using Grafen’s method (Grafen, 1989) implemented in the R package ‘ape’(Paradis, 2012).

Fixed effects were considered significant when 95% credible intervals did not overlap with 0 and pMCMC were less than 0.05 (pMCMC = percentage of iterations above or below a test value correcting for the finite sample size of posterior samples). Default fixed effect priors were used (independent normal priors with zero mean and large variance (10^10^)) and for random effects inverse gamma priors were used (V = 1, nu = 0.002). Each analysis was run for 1100000 iterations with a burn-in of 100000 and a thinning level of 1000. Convergence was checked by running each model three times and examining the overlap of traces, levels of autocorrelation, and testing with Gelman and Rubin’s convergence diagnostic (potential scale reduction factors <1.1).

#### Specific analyses

Two sets of analyses were conducted, one on the effect of relatedness on mean rates of mortality and pathogen abundances (*r*) and one on variances (*LnCVR*). All models were fitted with a Gaussian error distribution, study, species and phylogeny as random effects and each data point was weighted by the inverse sampling variance. Six analyses of mean effect sizes were conducted that had the following fixed effects (moderators): 1) intercept only model to test whether overall relatedness increased susceptibility to pathogens and increased mortality; 2) three-level factor of whether mortality was measured in the presence of pathogens, mortality was measured in the absence of pathogens, or whether the abundance of pathogens was examined (referred to here as ‘fitness measure’); 3) four-level factor of the effect of presence and absence of pathogens in experimental versus observational studies; 4) four-level factor of the effect of experimentally manipulating or observing relatedness in the presence and absence of pathogens; 5) eight-level factor of the effect of living with kin and non-kin in the presence and absence of pathogens in experimental and observational studies. All analyses were repeated for *LnCVR* apart from five, as variance estimates were only available for 7 species that live with non-kin.

#### Verification analyses

We checked the robustness of our results to potential inbreeding effects (Zr and LnCVR: Tables S14 & S15), whether studies were conducted in laboratories or under natural conditions (Zr & LnCVR: Tables S16 & S17), and the type of statistical analyses used in studies (Zr: Table S18). To check for any effects of inbreeding we repeated analysis four (see above) removing data points where inbreeding could potentially influence effect sizes (see Table S1 for effect size details. See Tables S14-16 for re-analysis). There was a large overlap in whether studies were conducted in laboratories and whether they were observational or experimental: All studies conducted in laboratories were experimental whereas observational studies had 141 effect sizes that were from the field and 23 from laboratory studies. To check for laboratory effects we therefore restricted data to observational studies and tested if effect sizes differed between laboratory and field studies (Tables S16 & 17). To examine the infleunce of the type of statistical tests used in studies (number of different analysis techniques: 15) we included ‘analysis technique’ as a random effect in our main model (analysis four above: see R script “ZrModels” M9). In all verification analyses our results remained unchanged and quantitatively similar (Tables S14-S18).

#### Testing for publication bias

Publication bias across studies was checked using funnel plot visualisation and Egger’s regression (Egger et al., 1997). Egger’s regressions of both Zr and LnCVR were performed by including the inverse sampling variance as a covariate in our full model (analysis 4 above: see R script “PublicationBias”). In both analyses, the slope of the inverse sampling variance was not significantly different from zero (BPMM: inverse sampling variance on Zr CI = −0.03 to 0.01 and LnCVR CI = −0.04 to 0.12) and funnel plots of residuals were also generally symmetrical (Figure S4). This suggests there is little evidence of publication bias.

## Acknowledgments

This research was funded by the Knut and Alice Wallenberg Foundation (Wallenberg Academy fellowship number 2018.0138) and the Swedish Research Council (grant number 2017-03880). We are very grateful to Dan Noble for comments on the manuscript.

## Competing interests

The authors have no competing interests.

## Author contributions

CKC, EO and HMB planned, designed and collected the data for the study. CKC performed the statistical analyses and wrote the paper with input from all authors.

## Data availability

All data, code and supplementary information are available at the open science framework (OSF): DOI 10.17605/OSF.IO/Q3ANE

